# Graph neural networks for integrated information and major complex estimation

**DOI:** 10.1101/2024.12.31.630856

**Authors:** Tadaaki Hosaka

## Abstract

This study investigates the potential of graph neural networks (GNNs) for estimating the system-level integrated information and major complex in integrated information theory (IIT) 3.0. Owing to the hierarchical complexity of IIT 3.0, tasks such as calculating integrated information and identifying major complex are computationally prohibitive for large systems, thereby restricting the applicability of IIT 3.0 to small systems. To overcome this difficulty, we propose a GNN model with transformer convolutional layers characterized by multi-head attention mechanisms for estimating the major complex and its integrated information. In our approach, exact solutions for integrated information and major complex are obtained for systems with 5, 6, and 7 nodes, and two evaluations are conducted: (1) a non-extrapolative setting in which the model is trained and tested on a mixture of systems with 5, 6, and 7 nodes, and (2) an extrapolative setting in which systems with 5 and 6 nodes are used for training and systems with 7 nodes are used for testing. The results indicate that the estimation performance in the extrapolative setting remains comparable to that in the non-extrapolative setting, showing no significant degradation. In an additional experiment, the model is trained on systems with 5, 6, and 7 nodes and tested on a larger system of 100 nodes, composed of two subsystems of 50 nodes each, with limited inter-subsystem connectivity resembling a split-brain configuration. When the connectivity between the subsystems is low, “local integration” emerges, meaning that a single subsystem forms a major complex. As the connectivity increases, local integration rapidly disappears, and the integrated information gradually rises toward “global integration,” in which a large portion of the entire system forms a major complex. Overall, our findings suggest that GNNs can potentially be used for estimating integrated information, major complex, and other IIT-related quantities.

**Author summary:** Understanding consciousness is one of the most profound challenges in science. Integrated information theory (IIT) provides a solution to this problem by assessing how information is intrinsically unified within a system, suggesting that consciousness emerges from this integrated structure. However, calculating the extent of this integration in large systems, e.g., the brain, is an extraordinarily complex task; such a calculation is currently feasible only in small systems. Our study explores how graph neural networks (GNNs), a type of deep learning model designed for analyzing graph structures, can estimate IIT-related measures without the intricate calculations required by IIT. By training GNNs on small-sized systems and testing them on more complex configurations, we find that GNN models are a promising candidate for approximating the key measures of information integration and identifying regions of high integration, known as “major complex,” which is considered essential for consciousness. This study opens possibilities for using artificial intelligence to model larger brain-like networks, potentially offering new insights into the nature of consciousness and the inner workings of complex systems.

## Introduction

Integrated information theory (IIT), proposed by Giulio Tononi, has witnessed significant development over the last two decades, evolving through four major versions [1–7] and becoming a central framework for understanding consciousness. IIT asserts that consciousness emerging in a system is equivalent to the integrated information quantified by Φ, which measures the information in the whole exceeding the sum of the information in its parts. This core principle is a fundamental concept shared across all versions of IIT.

IIT 1.0 [1] introduced the foundational notion that consciousness arises from the integrated information within a system, emphasizing the difference between the whole and its parts. In IIT 2.0 [2–4], this idea was rigorously formalized through the introduction of the “minimum information partition,” which is the division minimizing the decrease in the degree of information integration, compared to the non-partitioned system. However, IIT 2.0 treated the entire system as a single entity for calculating the integrated information without considering that consciousness might be composed of integrated subsystems or exploring any hierarchical structures within the system. By contrast, IIT 3.0 [5, 6] introduced a hierarchical approach that analyzes all possible subsystems to elaborate the nature of consciousness more precisely (S1 Text). For every group of nodes within each subsystem, the mechanism-level integrated information, denoted as *φ*, is measured according to the principle of the minimum information partition. From the collection of these values of *φ* and their associated probability distributions, the system-level integrated information for each subsystem is then computed as Φ. Finally, the subsystem with the highest Φ across the entire system is identified as the “major complex,” which represents the location of consciousness within the system. The latest version, IIT 4.0 [7], has significantly advanced the theory by introducing the concept of “relations” between node groups to refine how the value of Φ is derived from the collection of *φ*.

This study is based on IIT 3.0, whose primary objective is to identify the “major complex,” the subset of nodes with the highest integrated information Φ, given a system consisting of nodes and edges, node states, and state transition probabilities. The most notable advancement from IIT 2.0 to IIT 3.0 is the introduction of the aforementioned hierarchical structure to compute the system-level integrated information, which significantly increases the computational complexity owing to more intricate hierarchical procedures. Deriving integrated information Φ and major complex pose substantial computational challenges, not only because finding the minimum information partition requires evaluating all possible partitions of a target system but also because IIT 3.0 involves multiple nested combinatorial optimization problems, as described in S1 Text. Consequently, the exponential growth in computational complexity with the number of nodes restricts rigorous IIT 3.0 calculations to extremely small systems, typically those comprising fewer than 10 nodes, thereby limiting the applicability of IIT 3.0 to large-scale systems.

Several theoretical and numerical studies have explored the specific characteristics and limitations of IIT 3.0 [8–11]; however, practical approximation methods have not been proposed thus far. Although some approximation methods have been developed to efficiently identify optimal partitioning patterns [12, 13], they are applicable to the framework of only IIT 2.0 and not IIT 3.0, owing to the complex hierarchical structure. One exception is the “cut one” approximation [14], which limits the search for the minimum information partition to those that isolate a single node. However, this approximation method is clearly too crude and limited in its applicability, and it cannot fully address the computational difficulties posed by IIT 3.0.

To extend the applicability of IIT to larger systems, it is crucial to address its computational challenges. Graph neural networks (GNNs) offer a promising solution to this problem. GNNs are a type of deep learning model designed specifically for graph-structured data, where nodes represent entities and edges represent their relationships. GNNs have been effectively employed in various biological applications, including drug discovery [15], predicting protein interfaces [16, 17], modeling neural connectivity patterns in the brain [18], identifying relationships between diseases and genes [19], and classifying lung cancer subtypes from whole-slide images [20]. Although GNNs face challenges in terms of interpretability, they often outperform traditional machine learning methods by accurately extracting complex patterns from data. This could make GNNs particularly suitable for estimating integrated information and major complex in IIT 3.0, without explicitly modeling the intricate computational processes.

This study aims to explore the potential of GNNs as an approximation method for estimating the major complex and its maximum integrated information Φ. First, we evaluate the performance of GNNs on small-scale systems where exact theoretical calculations are feasible. After confirming reasonable accuracy in these small systems, we expand our experiments to larger systems that resemble a split-brain scenario [21, 22], consisting of two subsystems, in order to further assess the capability of GNNs in more distinct settings. Through these experiments, we demonstrate the potential effectiveness of GNNs as a practical approximation algorithm for IIT 3.0. Hereafter, the term “integrated information” and the variable Φ will refer to the maximum value of the system-level integrated information across all subsystems, i.e., the integrated information of the major complex, unless otherwise noted.

## Methods

In this study, we estimate integrated information Φ and major complex for randomly generated systems consisting of *N* (= 5, 6, 7) nodes. This limitation on the number of nodes is necessary because obtaining the exact solutions for IIT 3.0 becomes computationally infeasible for larger systems. For each value of *N*, numerous random systems are generated and used to train the GNN model.

Each node in the system can be in one of two states, +1 or −1, denoted as *S*_*i*_ (*i* = 1, 2, …, *N*), and the states are randomly assigned. The undirected edges between any pair of nodes are assigned with a probability of *p* = 0.4, and the edge weights *J*_*ij*_ between nodes *i* and *j* are randomly determined following a standard normal distribution. The conditional probability *p*_*i*_(*s*|***S***) that node *i* will take on state *s* ∈ {1, −1} in the next time step is determined on the basis of the current states ***S*** = (*S*_1_, *S*_2_, …, *S*_*N*_) using the Boltzmann distribution:

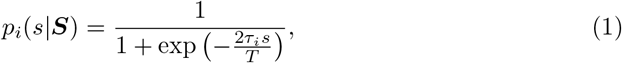

where *τ*_*i*_ = Σ_*j*∈𝒩 (*i*)_ *J*_*ij*_*S*_*j*_ represents the input to node *i*, and 𝒩 (*i*) denotes the set of nodes connected with node *i*. The parameter *T*, controlling the sharpness of the distribution, is uniformly sampled from the range [0.1, 3.0] and is set independently for each generated system. This stochastic model captures the dynamics of the system, allowing for the probabilistic determination of the state of each node on the basis of interactions with its neighbors.

Using the PyPhi library published by Mayner et al. [14], the exact solutions for the integrated information Φ and the corresponding major complex are obtained for each of the randomly generated systems. These systems and their solutions serve as the dataset for subsequent analysis within a supervised learning framework, where a GNN is trained on part of the dataset and tested on the remaining data.

Each system is treated as a graph where nodes and edges have specific features. Node *i* (*i* = 1, 2, …, *N*) is assigned a feature vector consisting of the following six dimensions:

1. max {*p*_*i*_(1|***S***), *p*_*i*_(−1|***S***)} : The larger value between the probability that node *i* will be in state +1 in the next time step and the probability that it will be in state −1. Note that the node states ***S*** affect only this feature and have no influence on the other features.
2. Parameter *T* : A parameter that controls the sharpness of the probability distribution. In each graph, all nodes share the same value of *T*.
3. Node degree: The number of edges connected to node *i*, indicating its connectivity in the system.
4. Closeness centrality: A measure of how close node *i* is to all other nodes in the system, calculated as the inverse of the average of the shortest path distances from node *i* to all other nodes. It indicates how rapidly information spreads from node *i* to others.
5. Betweenness centrality: A measure of how frequently node *i* lies on the shortest paths between pairs of other nodes. It indicates the importance of node *i* in the network communication.
6. Clustering coefficient: A measure of how closely the neighbors of node *i* are connected to each other, indicating the density of local connections around node *i*. It is calculated as the ratio of actual connections to possible connections among the neighbors of node *i*.

Meanwhile, the edge features simply consist of a single dimension representing the edge weights, *J*_*ij*_, between the nodes.

It should be noted that the node state *S*_*i*_ is not utilized as a feature. The random systems in our study exhibit a high degree of symmetry with respect to the inversion of the state vector. Specifically, the conditional probabilities of transitions are symmetric, indicating that *p*(***S***^*^|***S***) = *p*(−***S***^*^ |−***S***), where ***S***^*^ denotes a specific state to which the system transitions from the current state ***S***. These conditional probabilities are derived by the product of Eq 1 over all the nodes, assuming conditional independence.

According to IIT 3.0, for such symmetrical systems, both the state vector ***S*** and its inverse −***S*** should yield the same integrated information and major complex. However, if the node state *S*_*i*_ is included as a node feature in the GNN, the use of ReLU activations and bias terms could disrupt this symmetry, leading to different outputs for ***S*** and − ***S*** despite their theoretical equivalence. We have designed the six aforementioned features to be identical for both node states ***S*** and −***S***, ensuring that the GNN produces theoretically consistent predictions.

Table 1 shows our GNN architecture, which is designed to perform multi-task estimation, i.e., predicting the integrated information Φ and identifying the major complex for a target system. The input consists of the six-dimensional feature vector for every node, with each dimension normalized using the entire training dataset. These inputs pass through four consecutive transformer convolutional layers, where attention weights are dynamically adjusted on the basis of the relationships between nodes [23]—a mechanism that has achieved significant success in natural language processing (NLP) [24]. In addition, our model employs the multi-head attention mechanism, where each head is expected to independently capture different aspects of the relationships between nodes. Specifically, *d*-dimensional feature vector of each node, ***x***_*i*_ ∈ *ℝ*^*d*^, is transformed into three components, namely query (***q***_*i*_), key (***k***_*i*_), and value (***v***_*i*_), through learnable weight matrices ***W*** _*q*_, ***W*** _*k*_, and ***W*** _*v*_. The transformations for each head *h*(= 1, 2, …, *H*) are given by

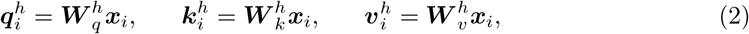

where 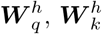, and 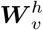 are in ℝ^*C×d*^ and *C* represents the number of output channels. The attention weight toward node *i* from its neighbor *j* is computed using the dot product of the query from node *i* and the key from node *j*, with the addition of a term based on the edge feature ***e***_*ij*_ (one dimension in our study) and its learnable transformation matrix 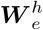:

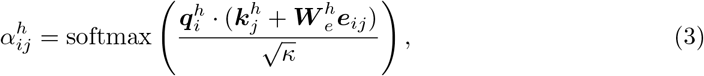

where 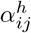 represents the importance of the feature of node *j* for updating node *i*, and *κ* is a scaling parameter typically set to *C/H*. The updated feature for node *i* in each head is then computed as a weighted sum of the value vectors from its neighboring nodes:

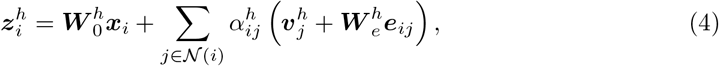

where matrix 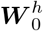 is in ℝ^*C×d*^. The outputs from the multiple heads are concatenated to form the final updated feature for node *i*:

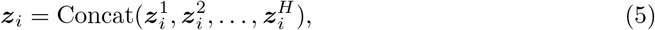

which results in a feature vector ***z***_*i*_ of dimension *CH*. This multi-head attention mechanism enables our GNN to attend to different aspects of the input graph simultaneously. Furthermore, batch normalization and dropout are applied to the first three transformer convolutional layers in order to stabilize training and prevent overfitting.

**Table 1.** Network architecture of the GNN for estimating the integrated information and the major complex.

| Layer | Description |
| --- | --- |
| Input | Normalized features (Six-dimensional) |
| Transformer Convolution 1 | 50 output channels, 4 heads |
| Batch Normalization | ReLU activation |
| Dropout | Drop rate of 0.3 |
| Transformer Convolution 2 | 150 output channels, 4 heads |
| Batch Normalization | ReLU activation |
| Dropout | Drop rate of 0.3 |
| Transformer Convolution 3 | 150 output channels, 4 heads |
| Batch Normalization | ReLU activation |
| Dropout | Drop rate of 0.3 |
| Transformer Convolution 4 | 50 output channels, one head |
| Split to Branch 1 and Branch 2 |  |
| **[Branch 1]** |  |
| Global Max Pooling |  |
| Dropout | Drop rate of 0.3 |
| Full Connection 1 | Linear activation, one output corresponding to the value of integrated information $\Phi$ |
| **[Branch 2]** |  |
| Global Max Pooling Difference | Subtraction of the global max-pooled vector from the feature vector of each node |
| Dropout | Drop rate of 0.3 |
| Full Connection 2 | Softmax activation, two outputs for major complex classification |

Following the fourth transformer layer, the network splits into two separate branches. In the first branch, the graph-level features are obtained using global max pooling, which selects the most prominent node features across the entire graph. Then, this pooled feature vector is passed through a dropout layer, followed by a fully connected layer that outputs the estimated integrated information Φ for the entire system.

In the second branch, the global max-pooled vector is subtracted from the feature vector of each node. This operation highlights the relative importance of each node compared to the most prominent features across the graph. Subsequently, the resulting differences are passed through a fully connected layer with a softmax activation function to perform a binary classification, determining whether each node is included in the major complex.

The subtraction of the global max-pooled vector from the feature vector of each node is crucial for improving the classification performance. Instead of this subtraction, relying solely on individual node features as inputs to the fully connected layer in Branch 2—an approach commonly used in node classification tasks—can cause significant errors when dealing with systems composed of multiple disconnected subsystems, such as subsystems A and B, where no edges exist between them. As illustrated in Fig 1(a), we assume that the GNN accurately predicts the integrated information with Φ_*A*_ *>* Φ_*B*_ and identifies the major complex for each subsystem when they are evaluated independently, i.e., system A or system B alone constitutes the whole system. Even under this assumption, the prediction fails when subsystems A and B are evaluated together as a single disconnected larger system (Fig 1(b, c)). In this larger system, the graph convolution results for each subsystem A or B remain identical to those when they are evaluated individually, causing the GNN to mistakenly classify some nodes from the lower integrated information subsystem (B in this case) as part of the major complex. This misclassification occurs because using only the individual node features without considering the global structure fails to differentiate between the subsystems. By subtracting the global max-pooled vector from the feature vector of each node, the model takes into account the relative importance of each node, increasing the likelihood that only the nodes from the subsystem with the highest integrated information (A in this case) are correctly classified as part of the major complex.

**Fig 1.**
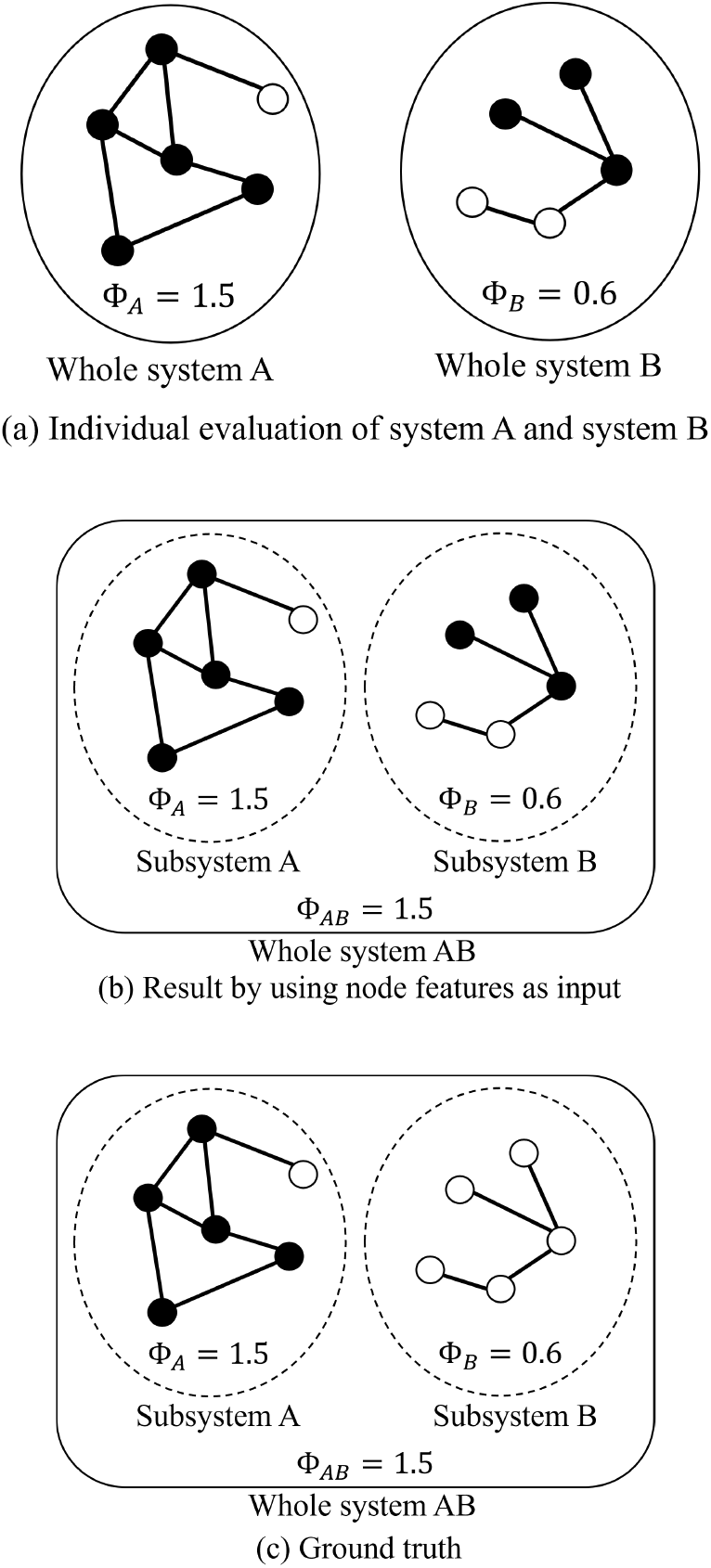
Integrated information Φ and major complex in a system consisting of two disconnected subsystems. (a) When each system is evaluated individually, integrated information and major complex can be correctly estimated using node features alone, without subtracting the max-pooled vector as input to the fully connected layer. Black circles represent nodes included in the major complex, while white circles represent nodes not included. (b) Since no edge exists between the two subsystems, the results of the transformer convolutions are identical to those obtained when each subsystem is evaluated independently. However, this leads to an incorrect estimation. (c) In the ground truth, the nodes in the subsystem with the smaller Φ should not be included in the major complex.

## Results

In this section, we present the results of our simulations. We employed the PyTorch Geometric library [25] to implement the GNNs in the following experiments. To evaluate the performance of our GNN model in estimating integrated information Φ and classifying the inclusion of the nodes in the major complex, we conducted three types of experiments.

The first experiment used a dataset of 3000 graphs in total, consisting of 1000 random systems generated as described in the previous section for each case of *N* = 5, 6, and 7. The entire dataset was randomly shuffled and then split into two subsets: 90% of the total data was used in the training process, while the remaining 10% was reserved for testing. Then, the performance of the model was evaluated on the test dataset by measuring the accuracy of both the integrated information estimation and the classification of whether each node belongs to the major complex.

The second experiment aimed to assess the ability of the model to generalize to larger systems than those used in training, which was regarded as an extrapolative setting. We trained the model using a dataset consisting of 1500 graphs for each case of *N* = 5 and 6, i.e., a total of 3000 random systems. The test dataset consisted of 1000 random systems for *N* = 7. The performance of the model was evaluated on this test dataset with the same metrics as in the first experiment.

Finally, we predicted integrated information and major complex for larger systems of *N* = 100 using the GNN models trained in the first experiment. Exact solutions for such large systems are computationally infeasible. To facilitate qualitative interpretation, we designed a test system like a split brain [21, 22], consisting of two subsystems, each containing 50 nodes. By varying the probability of an edge existing between nodes in different subsystems, we investigated the changes in the estimated integrated information and major complex, highlighting the transition from local to global integration.

For the three types of experiments, we applied several common settings and parameters that were empirically determined. We defined two class labels: label *in*, representing nodes “included in the major complex,” and label *out*, representing nodes “not included in the major complex.” To address the imbalance between the numbers of these classes, meaning that label *in* was more frequent in our dataset, we applied a penalty factor of 1.8 to label *out* in the cross-entropy calculation. The total loss function for our multi-task model consisted of the sum of the mean squared error for the integrated information estimation and five times the cross-entropy loss for the major complex classification. This weighting factor of five was empirically introduced to balance the two tasks. The optimizer was set to LION (evolved sign momentum) [26], which updates parameters solely on the basis of the sign of the gradients, in contrast to traditional optimizers like SGD or Adam [27], which consider the magnitude of the gradients. The LION optimizer provides a simple yet effective optimization strategy, focusing on efficiency and memory-saving features suitable for training deep network models. The learning rate in the algorithm was set to 0.0001 and the mini-batch size was set to 128.

Furthermore, to address the imbalance between label *in* and label *out* in the training dataset, we adopted a data augmentation strategy to artificially increase the number of label *out* instances. Specifically, we generated additional systems amounting to 5% of the total data used in the training process by randomly selecting two graphs from the training dataset and treating them as disconnected subsystems within a larger system. Assuming that the integrated information values of the two selected graphs are Φ_*A*_ and Φ_*B*_ with Φ_*A*_ *>* Φ_*B*_ as in Fig 1(a), the whole of this artificially created system would have an integrated information value of Φ_*A*_. The true label for major complex inclusion would be retained for nodes from graph A, while all nodes from graph B would be assigned the label *out* as in Fig 1(c), thereby increasing the proportion of *out* labels in the dataset.

As the distribution of Φ values within the training dataset was not uniform, an oversampling technique was adopted to improve the estimation accuracy in the low-frequency regions. We used k-means clustering to divide the data into seven bins based on the value of Φ. Then, we oversampled the smaller bins to match the size of the largest bin, adding noise by applying a multiplicative factor of 5% independently to both the integrated information values Φ and each dimension of the node features.

Following the application of data augmentation and oversampling techniques, 10% of the expanded training dataset was separated and utilized as validation data to monitor the performance of the model during training. The training process was halted if the validation loss did not improve for 50 consecutive epochs, following the early stopping strategy.

### Estimation in non-extrapolative setting

The results of the first experiment are summarized in Table 2. The table includes four main metrics evaluated on the test data: the mean squared error (MSE) and correlation coefficient between the estimated and actual values of the integrated information Φ, the accuracy of the node-wise prediction for inclusion in the major complex, and the graph-wise accuracy, i.e., the proportion of graphs where the major complex is predicted completely correctly. The values shown in the table are averages obtained over 100 repetitions of the experiment, where different combinations of training and testing datasets were used. The table presents the performance achieved with the proposed method, and for comparison, it also shows the following:

**Table 2.** Performance evaluation with consistent training and testing node sizes (non-extrapolative). The performance of the proposed method is compared against variations in node features, convolution types, optimizers, and pooling strategies, as well as single-task estimations. The numbers in parentheses for the proposed method indicate its rank in each column, and results outperforming the proposed method are underlined.

| Category | Configuration | Integrated information estimation |  | Major complex estimation |  |
| --- | --- | --- | --- | --- | --- |
|  |  | MSE | Correlation | Node-wise acc. | Graph-wise acc. |
| Proposed method |  | 0.4611 (2) | 0.7446 (2) | 0.8574 (2) | 0.5779 (3) |
| Removed feature | Probability | 0.5272 | 0.7061 | 0.8544 | 0.5615 |
| | Parameter $T$ | 0.5128 | 0.7149 | 0.8492 | 0.5566 |
|  | Node degree | 0.5122 | 0.7076 | 0.8554 | 0.5727 |
|  | Closeness centrality | 0.5585 | 0.6870 | 0.8521 | 0.5663 |
|  | Betweenness centrality | 0.4920 | 0.7206 | 0.8550 | 0.5680 |
|  | Clustering coefficient | 0.5362 | 0.7061 | 0.8443 | 0.5437 |
| Substitute convolution | GraphConv | 0.6254 | 0.6799 | 0.8107 | 0.4650 |
|  | GAT | 0.5706 | 0.6950 | 0.7865 | 0.4168 |
| Substitute optimizer | Adam | 0.4828 | 0.7331 | 0.8572 | <u>0.5780</u> |
|  | RAdam | 0.4899 | 0.7364 | <u>0.8589</u> | <u>0.5793</u> |
|  | AdamW | 0.4780 | 0.7413 | 0.8565 | 0.5742 |
| Single task | $\Phi$ estimation | 0.4638 | 0.7376 | - | - |
|  | Major complex estimation | - | - | 0.8557 | 0.5755 |
| Pooling | Global average | 0.4712 | 0.7380 | 0.8397 | 0.5207 |
|  | Not subtraction | <u>0.4495</u> | <u>0.7547</u> | 0.8281 | 0.4908 |

- Performance when removing each dimension of the node features.
- Performance when replacing the transformer convolutional layer with more standard convolutional types, specifically the graph convolutional layer (GraphConv class provided in the PyTorch Geometric library) and graph attention network [28] (GAT class in the PyTorch Geometric library). GraphConv simply aggregates features from neighboring nodes using a weighted sum without attention mechanisms. GAT employs simpler attention mechanisms than the transformer convolution.
- Performance when replacing LION with other optimizers, specifically Adam [27], which is widely regarded as the de facto standard, and its derivatives RAdam [29] and AdamW [30].
- Performance for single-task scenarios, where either integrated information Φ or major complex is estimated independently. This corresponds to evaluations on networks with either Branch 1 or Branch 2 removed.
- Performance when using global average pooling instead of global max pooling in Branches 1 and 2, as well as when using the node features without subtracting the max-pooled vectors as inputs to the fully connected layer in Branch 2.

The performance of the proposed method in estimating integrated information shows a relatively strong correlation coefficient of 0.7446 between the estimated and true values, implying a reasonable level of predictive capability. Fig 2 shows a scatter plot comparing the estimated and actual values of Φ in one trial. Most systems had the values of Φ below 1, where the model demonstrated higher accuracy. By contrast, systems with Φ values around 1 or higher were less frequent and tended to be predicted with lower accuracy. The uneven distribution of samples in the dataset, with fewer samples for higher Φ values, leaded to greater variability in predictions for those regions. While oversampling could increase the apparent number of samples, it did not sufficiently reproduce the diverse characteristics of graph structures and node features in regions with few samples, which might limit its effectiveness in improving prediction accuracy. This result suggests that having a more balanced set of real samples could help improve the model’s robustness across the entire range of Φ values.

**Fig 2.**
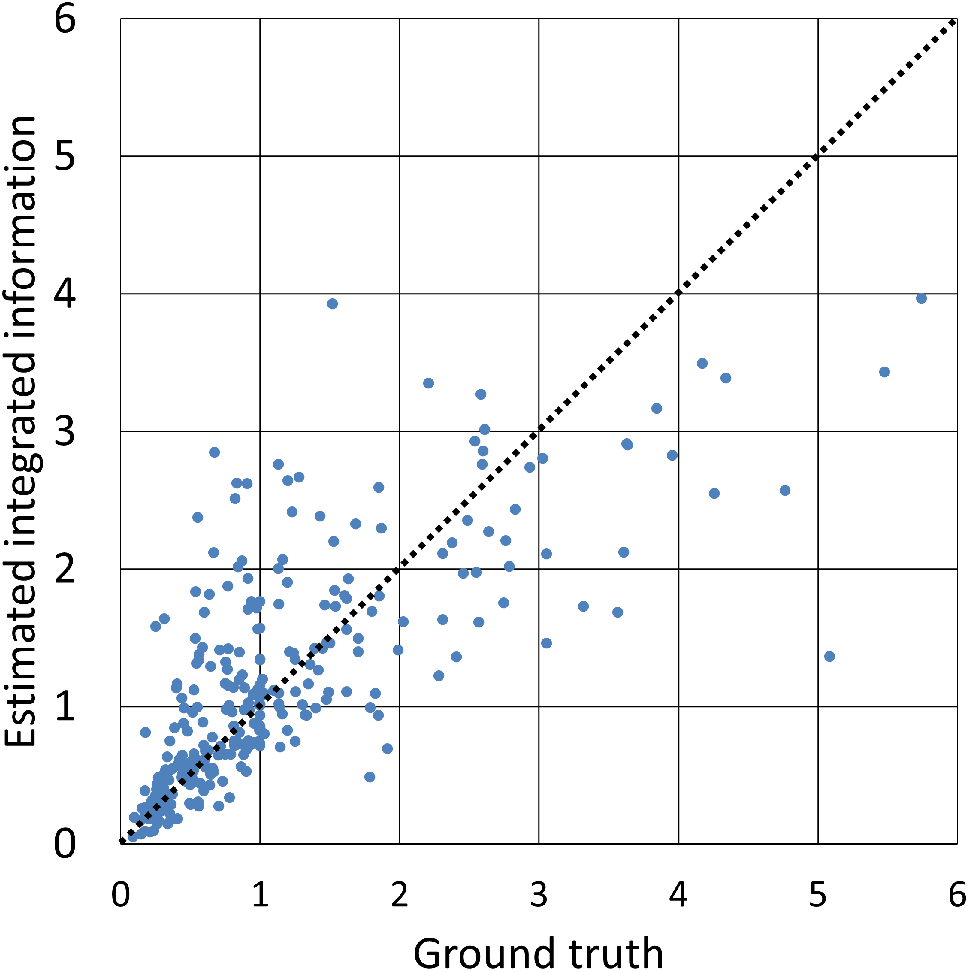
Scatter plot between estimated and true integrated information (non-extrapolative). All 300 test data used in one of the 100 trials conducted with the proposed method are shown. In this case, the correlation coefficient was 0.7464, which was close to the average over all the trials. Data for which the estimation is accurate are on the dotted line with a slope of 1.

When predicting node inclusion in the major complex, the proposed method achieves a node-wise accuracy of 0.8574, indicating reliable performance in this binary classification task. The graph-wise accuracy, which measures the proportion of graphs where the major complex is completely identified, is 0.5779. Interestingly, this is higher than the expected value obtained by simply raising the node-wise accuracy to the power of the number of nodes (5, 6, or 7). This suggests that our model captures a certain degree of interdependence among nodes forming the major complex, rather than treating each node independently. Specifically, the model seems to learn characteristic patterns in the graph structure associated with major complex, and to make incorrect predictions on graphs that deviate from these learned patterns. Further improvements can be achieved by intensively training on rare graph configurations.

When a single node feature is removed, the decline in accuracy for major complex estimation is relatively small. By contrast, the performance related to Φ estimation shows a more pronounced decrease, indicating that integrated information is more sensitive to changes in node features. It is intuitively expected that the higher probability of a node being in state +1 or −1 and the parameter *T*, both of which are closely related to the state transition probabilities required for the IIT calculation, would have a larger impact than the centrality-based features. However, we do not find significant differences that allow us to definitively determine the superiority of individual features. Meanwhile, the selection of convolution type has a greater influence on performance compared to node feature reduction. The standard convolutional methods, GraphConv and GAT, resulted in much lower performance compared to our transformer-based approach. This result highlights the effectiveness of the weight determination of the attention mechanism in the transformer, which has also proven successful in NLP.

In addition, although the Adam and the RAdam optimizers slightly outperform the proposed method in terms of major complex estimation accuracy, the overall performance of LION, Adam, RAdam, and AdamW shows no significant differences. However, the proposed method using LION demonstrates a clear advantage in efficiency, with an average convergence time of 94.2 epochs, compared to 166.2 epochs for Adam, 185.4 epochs for RAdam, and 160.8 epochs for AdamW. This highlights the effectiveness of the LION optimizer, designed primarily to enhance efficiency rather than accuracy, which is evident in its significantly faster convergence.

Estimating integrated information and major complex with the multi-task approach resulted in limited performance improvements compared to single-task models. This suggests that, although deriving integrated information and identifying major complex are theoretically related within the framework of IIT 3.0, the proposed method does not fully capture their deeper interconnections. Enhancing the ability of the multi-task model to better utilize the intricate relationships between integrated information and major complex is a key challenge for future work.

Using global average pooling, which is a standard choice for graph-wise estimation, fails to achieve the same level of accuracy as global max pooling, especially for major complex estimation. This highlights the importance of global max pooling in the proposed method. Global max pooling is more effective in capturing the influence of the most decisive or impactful node within the graph, which is necessary for distinguishing the major complex, whereas average pooling dilutes this influence and leads to reduced accuracy. Similarly, using node features directly as inputs to the fully connected layer in Branch 2, without subtracting the global max-pooled vector, significantly degrades the major complex estimation accuracy. Although this approach slightly improves estimation accuracy of Φ, subtracting the max-pooled vector, as mentioned earlier, offers a distinct theoretical advantage for major complex estimation, outweighing the minor gains in integrated information estimation.

### Estimation in extrapolative setting

The results of the second experiment, where the model was trained and validated on graphs with *N* = 5, 6 and tested on graphs with *N* = 7, are presented in Table 3. This setup represents an extrapolative scenario, as the model must generalize to larger graphs than those used during training. The structure of the table and metrics are consistent with those in the first experiment, and the values represent averages over 100 trials.

**Table 3.**
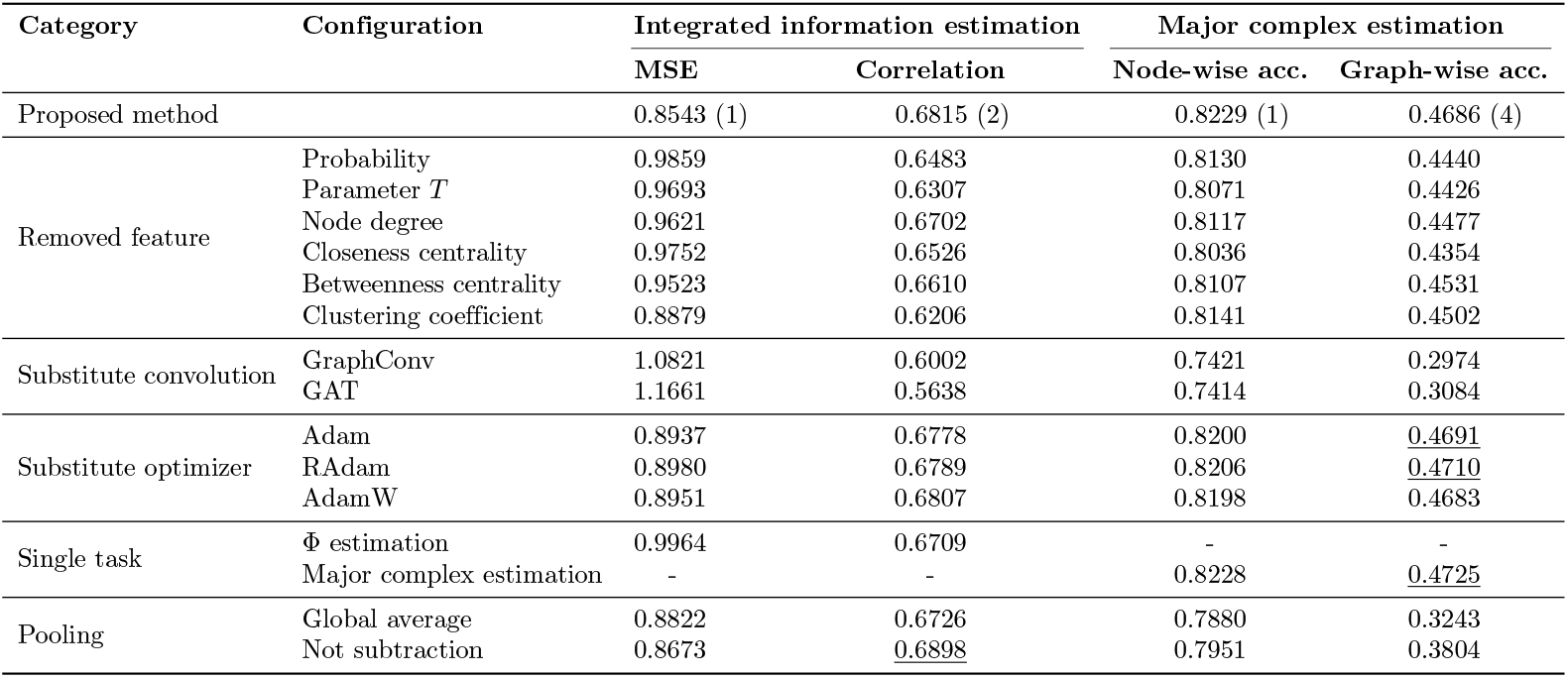
Performance evaluation with training on graphs for *N* = 5, 6 and testing on graphs for *N* = 7 (extrapolative). The same comparative experiments as in the previous experiment were conducted. The numbers in parentheses for the proposed method indicate its rank in each column, and results outperforming the proposed method are underlined.

| Category | Configuration | Integrated information estimation |  | Major complex estimation |  |
| --- | --- | --- | --- | --- | --- |
|  |  | MSE | Correlation | Node-wise acc. | Graph-wise acc. |
| Proposed method |  | 0.8543 (1) | 0.6815 (2) | 0.8229 (1) | 0.4686 (4) |
| Removed feature | Probability | 0.9859 | 0.6483 | 0.8130 | 0.4440 |
| | Parameter $T$ | 0.9693 | 0.6307 | 0.8071 | 0.4426 |
|  | Node degree | 0.9621 | 0.6702 | 0.8117 | 0.4477 |
|  | Closeness centrality | 0.9752 | 0.6526 | 0.8036 | 0.4354 |
|  | Betweenness centrality | 0.9523 | 0.6610 | 0.8107 | 0.4531 |
|  | Clustering coefficient | 0.8879 | 0.6206 | 0.8141 | 0.4502 |
| Substitute convolution | GraphConv | 1.0821 | 0.6002 | 0.7421 | 0.2974 |
|  | GAT | 1.1661 | 0.5638 | 0.7414 | 0.3084 |
| Substitute optimizer | Adam | 0.8937 | 0.6778 | 0.8200 | <u>0.4691</u> |
|  | RAdam | 0.8980 | 0.6789 | 0.8206 | <u>0.4710</u> |
|  | AdamW | 0.8951 | 0.6807 | 0.8198 | 0.4683 |
| Single task | $\Phi$ estimation | 0.9964 | 0.6709 | - | - |
|  | Major complex estimation | - | - | 0.8228 | <u>0.4725</u> |
| Pooling | Global average | 0.8822 | 0.6726 | 0.7880 | 0.3243 |
|  | Not subtraction | 0.8673 | <u>0.6898</u> | 0.7951 | 0.3804 |

For integrated information estimation, although the performance appears to decline compared to the previous experiment, the MSE and the correlation coefficient only for *N* = 7 in the first experiment were 0.7983 and 0.6730, respectively, which are comparable to those in this experiment. Fig 3 shows a scatter plot of estimated versus true integrated information values for a specific trial, illustrating the tendency of the model to underestimate Φ in the high-value range that is not covered by the training data. As higher integrated information values are associated with larger systems in most cases, the integrated information is expected to rise further as the graph size increases beyond *N* = 7; hence, the confidence in integrated information estimates may decrease for such larger graphs.

**Fig 3.**
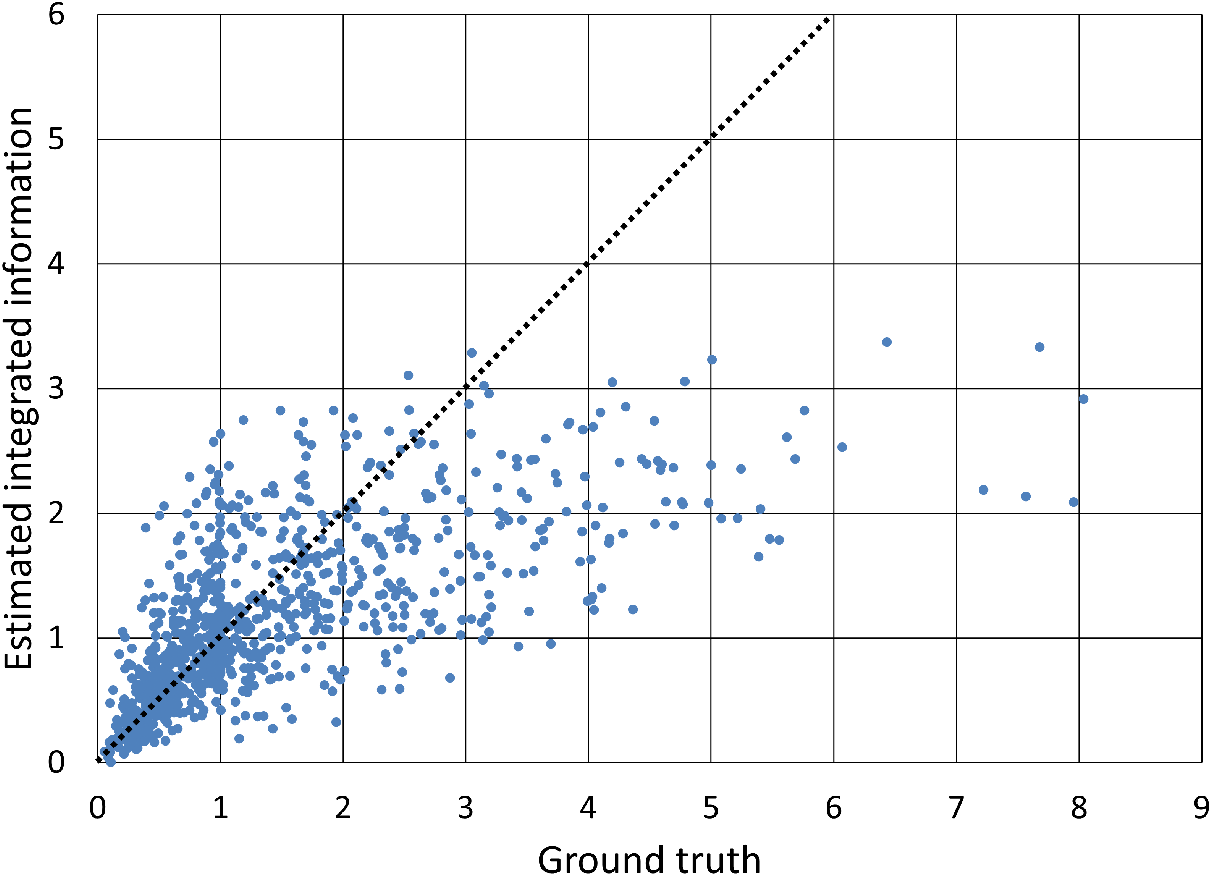
Scatter plot between estimated and true integrated information (extrapolative). The plot shows all 1000 test data corresponding to graphs for *N* = 7 in a specific trial conducted using the proposed method. In this trial, the correlation coefficient was 0.6815, matching the average across all trials.

For the major complex estimation task, both node-wise and graph-wise accuracies also appear to be lower compared to the first experiment. However, considering the test data only for *N* = 7 in the first experiment, the node-wise accuracy was 0.8302, and the graph-wise accuracy was 0.4893, which are nearly consistent with the results here. While the integrated information Φ behaves like an extensive variable and is sensitive to changes in graph size, the membership of each node in the major complex is inherently non-extensive. This suggests that our model may maintain a certain level of reliability in predicting the major complex for larger graphs and exhibit promising extrapolation performance.

In the comparison experiments, trends qualitatively similar to those observed in the previous experiment are found. The removal of one node feature leads to minor declines in major complex estimation accuracy, while integrated information estimation is more sensitive to these changes. The choice of convolution type also has a significant impact, with traditional methods resulting in lower performance than the transformer-based approach. The optimizer selection does not cause a significant difference in performance.

Our multi-task approach does not show a clear benefit over single-task estimation, suggesting potential limitations of the proposed method in fully leveraging the theoretical relationship between integrated information and major complex. In addition, global max pooling outperforms global average pooling especially in major complex estimation, as it captures the influence of the most significant node within the graph more effectively.

### Estimation for systems with larger nodes

This experiment aimed to evaluate the applicability of the proposed method to larger test systems by examining the behavior of estimated integrated information and major complex for systems of *N* = 100. Each test system comprised two subsystems: for the first 50 nodes, edges were randomly generated between pairs of nodes with a probability of 0.6, while the remaining 50 nodes were connected with a probability of 0.4. An edge connecting nodes between the two subsystems was set with probability *p*_*e*_ ranging from 0 to 0.4. For all edges, their strengths were independently sampled from a standard normal distribution. In the case of *p*_*e*_ = 0, the two subsystems are completely separate as in Fig 1, resembling a “split-brain” scenario. As the value of *p*_*e*_ increases, inter-subsystem connections are introduced, resulting in progressively greater integration between the subsystems. Specifically, we were interested in observing how the system transitions from the local integration to the global integration.

We generated 50 such large-scale test systems and applied 100 estimation models created in the first experiment of the non-extrapolative setting, which were trained on smaller systems with *N* = 5, 6, and 7, to predict the integrated information and identify the major complex for each test system. To enhance reliability, considering the variability in individual model predictions especially for extrapolative conditions, we averaged the estimated values of Φ across the 100 models and determined that a node belonged to the major complex if at least 60% of these models supported its inclusion.

Fig 4 shows the percentage of cases in which the subsystem of the first 50 nodes forms the major complex out of the 50 test systems by varying the value of *p*_*e*_. In all trials with *p*_*e*_ = 0, the major complex is composed of the first 50 nodes with denser internal connectivity. Identifying a single subsystem as the major complex is facilitated by the data augmentation techniques described in the Methods section and the operation of subtracting the max-pooled vector from the feature of each node. As *p*_*e*_ increases, this proportion of “local integration” declines rapidly, and when *p*_*e*_ reaches around 0.01 to 0.02, the major complex spans both subsystems with nearly 100% certainty.

**Fig 4.**
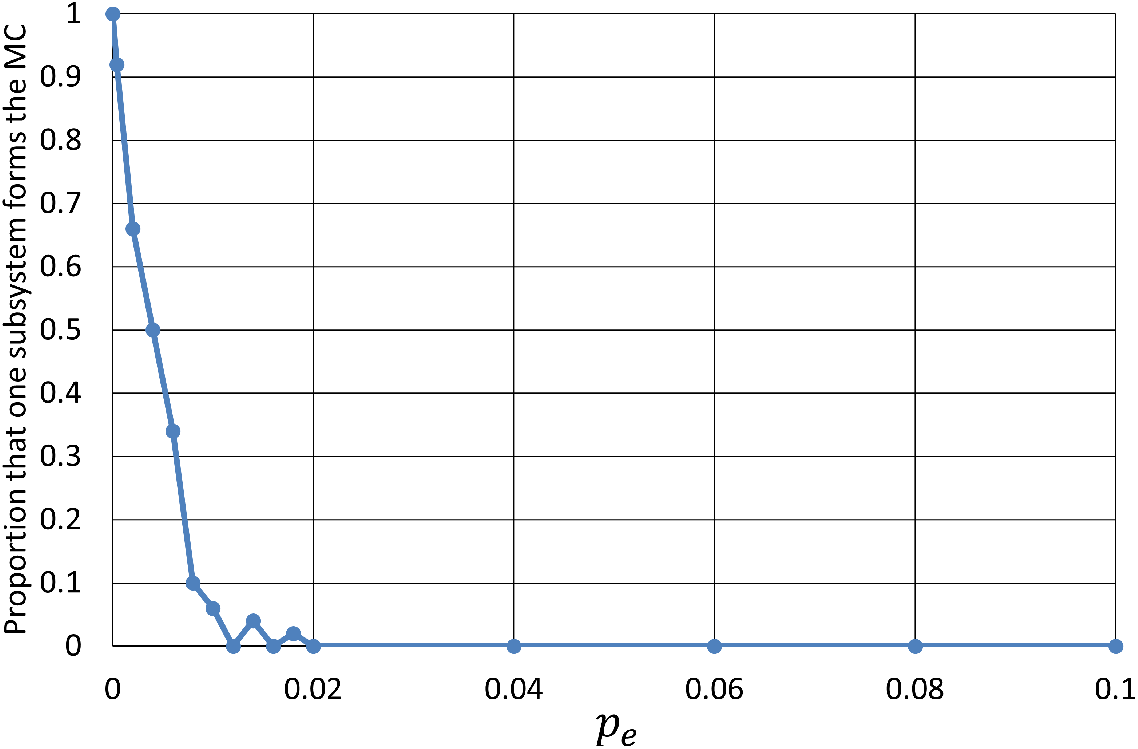
Proportion of cases out of 50 test systems where the single subsystem (the first 50 nodes) forms the major complex (MC) as a function of inter-subsystem edge probability *p*_*e*_. At *p*_*e*_ = 0, the major complex consists of only the first 50 nodes. As *p*_*e*_ increases, this rate rapidly declines.

Fig 5 shows the ratio of nodes included in the major complex out of the total 100 nodes as the value of *p*_*e*_ varies. This ratio was averaged over 50 test systems. At *p*_*e*_ = 0, the major complex is formed by all nodes in the first subsystem across all test systems, resulting in a ratio of 0.5. As *p*_*e*_ increases toward 0.02, the local integration disappears rapidly; by contrast, this ratio progresses more slowly, reaching only around 0.6. As *p*_*e*_ increases further, the ratio rises steadily and eventually reaches approximately 0.93, indicating that a larger fraction of the entire system contributes to the formation of the major complex, which can be regarded as “global integration.” This behavior may be akin to a functional shift from localized processing to a more distributed and unified form of information integration, as observed in neural circuits of the brain.

**Fig 5.**
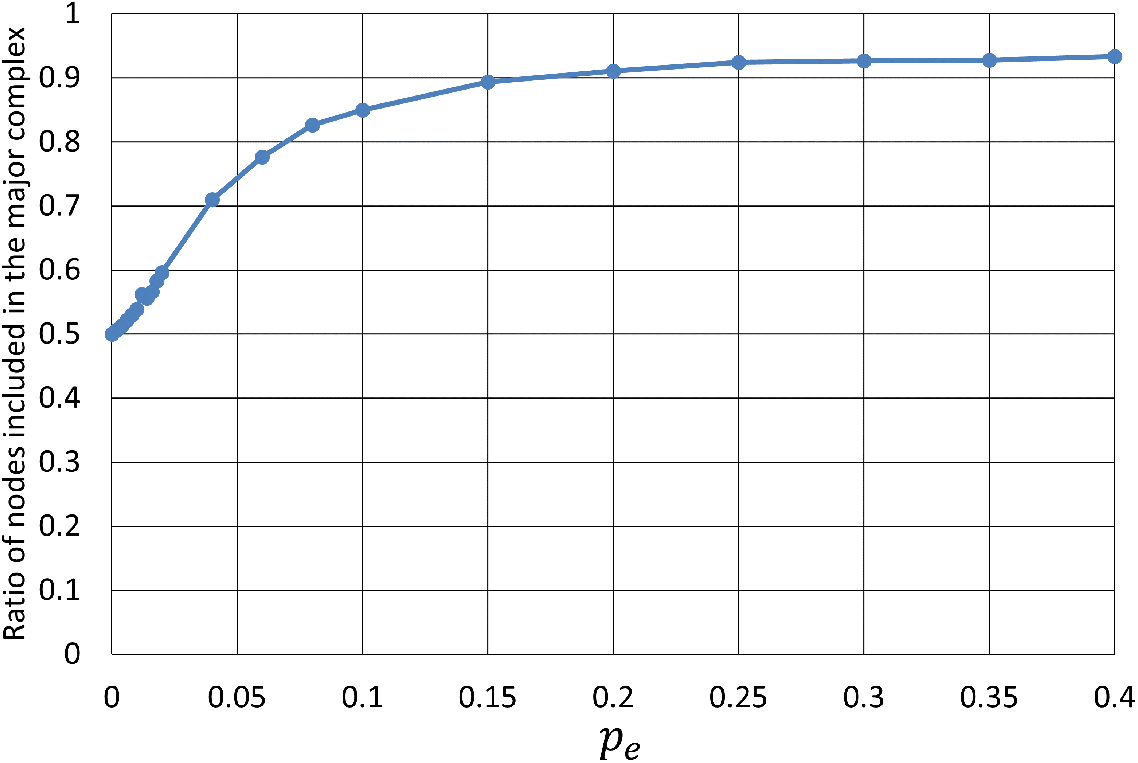
Ratio of nodes included in the major complex out of 100 total nodes as a function of *p*_*e*_. The values of ratios were averaged over 50 test systems. The ratio is 0.5 at *p*_*e*_ = 0, and it gradually increases with *p*_*e*_, eventually reaching around 0.93.

Fig 6 shows the estimated integrated information Φ averaged over 50 test systems and 100 models as a function of *p*_*e*_. Based on the results from previous experiments, the estimated values of Φ may not be highly reliable; however, relative changes in Φ can provide insights into significant trends. When *p*_*e*_ is sufficiently small (*p*_*e*_ ≲ 0.02) such that test systems resemble a split brain, the values of Φ remain approximately constant. As *p*_*e*_ increases and the major complex expands to include more nodes from both subsystems, the value of Φ grows correspondingly, indicating that enhanced interconnections and interactions between subsystems correlates with higher integrated information. As with the increase in the major complex size discussed above, such growth in the value of Φ might correspond to a transition from localized to cohesive and unified processing in the brain.

**Fig 6.**
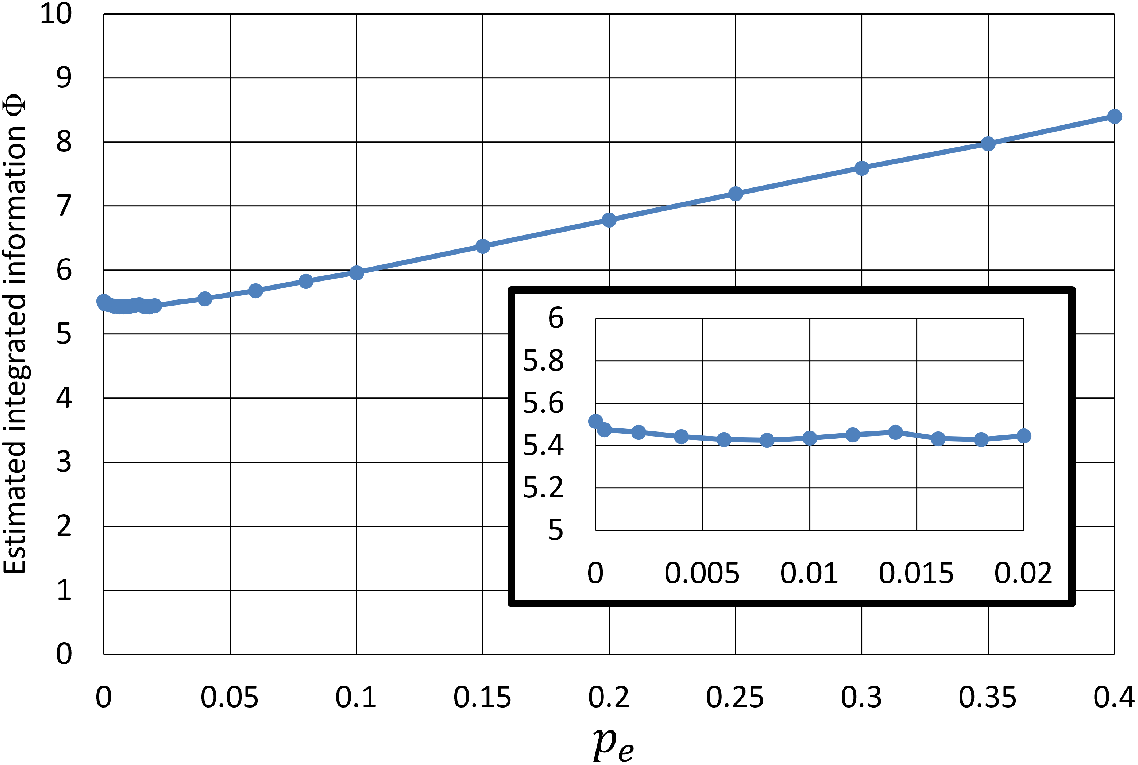
Estimated integrated information Φ as a function of *p*_*e*_. The values were averaged over 50 test systems and 100 estimation models. For small *p*_*e*_, the values of Φ remain nearly constant as illustrated in the interpolated diagram. It begins to increase as the major complex spans both subsystems, reflecting higher integration with increased connectivity.

These results illustrate the capacity of our model to differentiate between local and global integration in the system. For lower *p*_*e*_ values, the system exhibits characteristics similar to a split brain, where the major complex is restricted to one of the subsystems. As *p*_*e*_ increases and the inter-subsystem connectivity increases, the model begins identifying a unified major complex that spans both subsystems, corresponding to a globally integrated configuration. This highlights the potential of the GNN model to effectively make predictions based on large-scale connectivity patterns and to explore how system integration relates to neural configurations, although further validation across different datasets and scenarios is required.

## Discussion

This study explored the applicability of GNNs for estimating integrated information and major complex as defined by IIT 3.0. Our findings suggest that GNNs, especially those with multi-head transformer convolutional layers, can effectively capture relationships between nodes in small-scale systems and thus exhibit potential for extrapolation to larger, more complex systems. However, as a multi-task model, our GNN has limitations in fully capturing the intricate relationships between the integrated information and the major complex, suggesting that further refinement of the model architecture is necessary.

One potential enhancement involves expanding the predictive scope of the model to include additional variables (see S1 Text), such as the mechanism-level integrated information *φ* (small phi), which resides at a different level in the nested structure of IIT 3.0. Other relevant variables might include integrated information and major complex of unidirectionally-partitioned subsystems. As these variables span different computational stages within the IIT 3.0 framework, employing heterogeneous graphs [31, 32] to effectively link these distinct yet related variables could lead to more accurate estimation of the integrated information Φ and the major complex.

The proposed model faced challenges in accurately estimating integrated information as the number of nodes increased, particularly when extrapolating to larger systems. Underestimation in the region of higher values suggests that current training process might not sufficiently capture the scaling factors relative to the number of nodes. This issue could be addressed by treating the scaling factors as learnable parameters. One promising solution is to adopt an iterative learning approach. First, the network weights are trained on smaller graphs such as *N* = 5 or *N* = 6 with the scaling factors fixed. Next, only the scaling factors are trained on larger graphs such as *N* = 7 with the network weights fixed. By repeating these processes, the model may progressively enhance its extrapolative estimation ability.

Despite the aforementioned limitations of current models, the approach developed in this study can be applied to IIT 4.0, the latest version of the theory, because the main objective of IIT 4.0 remains the same: estimating integrated information Φ and identifying the corresponding complex (termed main complex in IIT 4.0) using the nodes, edges, and transition probabilities of a system as input. Furthermore, the advanced computational process introduced in IIT 4.0, including the concept of “relations” to specify connections between groups of nodes, could also be effectively represented by heterogeneous graph architectures.

From the formulation of the transformer convolution (Eq. 2), learnable parameters in the model are contained in the weight matrices ***W*** _*q*_, ***W*** _*k*_, ***W*** _*v*_, ***W*** _*e*_ and ***W*** _0_, which are shared across all nodes and are not directly influenced by the number of nodes *N*. By contrast, computing the attention coefficients *α*_*ij*_ (Eq. 3) is required for every edge in the graph. Additionally, deriving centrality-related node features becomes computationally expensive as *N* grows, since such computations often have a time complexity greater than *O*(*N*). Considering these factors comprehensively, if the number of connections per node remains *O*(1) and the centrality features are substituted with computationally efficient alternatives, the proposed method should remain feasible even for large *N* within the limits of available computational resources, particularly memory capacity.

In conclusion, although our GNN-based model is a promising approach for approximating IIT calculations, especially for large systems, further research is required to refine the architecture of the model, explore scaling strategies, and enhance the multi-task framework to leverage the strengths of each task for improving the prediction accuracy. Ultimately, developing such a tool will not only advance our ability to estimate integrated information and major complex in large-scale systems but also provide us with a deeper understanding of consciousness and other emergent phenomena, thereby offering insights for bridging computational models with neuroscience.

## Supporting information

**S1 Text. Nested optimization framework in IIT 3.0**

## Acknowledgments

This work was supported by JSPS KAKENHI Grant Number 23K11790.

